# Transcriptomic Analysis of Innate Immune Responses to *Toxoplasma gondii* in Bat Cells

**DOI:** 10.64898/2026.10.31.763326

**Authors:** Peter Yarovinsky, Samantha F. Kwok

**Author notes:** Correspondence: Peter Yarovinsky, Email.

## Abstract

Bats exhibit remarkable tolerance to numerous viral infections, a phenotype associated with distinctive adaptations in innate immune and inflammatory pathways. Whether these adaptations also shape responses to nonviral intracellular pathogens remains poorly understood. Here, we investigated the interaction between the protozoan parasite *Toxoplasma gondii* and epithelial cell lines derived from the black flying fox (Pteropus alecto). We found that both type I and type II strains of *T. gondii* readily invaded and replicated in bat cells, demonstrating a high degree of cellular permissiveness to infection. Surprisingly, transcriptomic analysis revealed remarkably limited early host responses despite progressive intracellular parasite replication. A pronounced transcriptional response emerged only at later stages of infection, coinciding with extensive parasite replication and cellular damage, and was characterized by changes in genes associated with inflammatory signaling, metabolic adaptation, and tissue remodeling. Together, these findings reveal an unusual interaction between bat cells and an intracellular protozoan pathogen, characterized by efficient parasite replication and markedly delayed host transcriptional activation. Our results suggest that distinctive regulation of cellular responses to intracellular infection may extend beyond viral pathogens and represent a broader feature of bat immune biology.

## Introduction

Bats (order Chiroptera) have emerged as a remarkable mammalian group because of their ability to harbor a wide range of viruses that cause severe disease in humans and other mammals while exhibiting little or no overt clinical illness themselves [1-3]. This unusual capacity has stimulated extensive investigation into bat immunity and has revealed multiple adaptations in antiviral innate immune pathways that promote effective control of viral replication while limiting excessive inflammatory responses [2-5]. Comparative studies have identified differences in interferon signaling, DNA sensing, inflammasome activation, and inflammatory cytokine production that are thought to contribute to the exceptional tolerance of bats to viral infection [2, 4-7]. These discoveries have positioned bats as an important model for understanding mechanisms that uncouple antimicrobial defense from immunopathology.

In contrast to the rapidly expanding knowledge of bat antiviral immunity, far less is known about how bats recognize and respond to bacterial, fungal, and parasitic pathogens. Whether the innate immune adaptations that underlie viral tolerance also influence host defense against these pathogens remains largely unexplored. This represents a significant gap in our understanding of bat immunobiology, particularly because innate immune recognition of nonviral pathogens relies on distinct pattern recognition receptors and signaling pathways that have received comparatively little attention in bats.

*T. gondii* is a ubiquitous intracellular protozoan parasite capable of infecting virtually all warm-blooded animals [8]. Host resistance to *T. gondii* depends critically on rapid innate immune recognition, IL-12 production, and the subsequent induction of IFN-γ, which together initiate antimicrobial mechanisms that restrict parasite replication [9-12]. Although the molecular pathways responsible for sensing *T. gondii* have been extensively characterized in mice and, to a lesser extent, in humans [13-15], it is currently unknown whether bats employ similar mechanisms or have evolved distinct strategies for detecting and controlling this parasite.

Epidemiological studies have demonstrated natural *T. gondii* infection in diverse bat species worldwide, indicating that bats are readily exposed to and can support infection by this parasite *in vivo* [16, 17]. Despite this evidence, virtually nothing is known about the cellular interactions between *T. gondii* and bat host cells or about the innate immune responses that are activated following infection. Understanding how bats recognize and control *T. gondii* may therefore provide new insights into the evolution of innate immunity against intracellular parasites and reveal whether the unique immunological adaptations identified in antiviral responses extend to protozoan infections.

To address these questions, we established an *in vitro* model of *T. gondii* infection using immortalized lung (PaLu) and kidney (PaKi) epithelial cell lines derived from the black flying fox (Pteropus alecto). We first examined the susceptibility of these cells to genetically distinct strains of *T. gondii* and their capacity to support intracellular parasite replication. We then performed transcriptomic analyses to characterize host responses to infection, focusing on innate immune signaling, inflammation, and cellular stress pathways. Together, these studies provide new insights into bat–protozoan interactions and explore whether the distinctive immune adaptations associated with antiviral defense in bats extend to nonviral intracellular pathogens.

## Results

### T. gondii efficiently infects and replicates in bat epithelial cells

Although epidemiological studies have provided evidence that bats can be naturally infected with *T. gondii* [16, 17], the ability of the parasite to directly infect and replicate in bat cells has not been well established. To address this question, we examined *T. gondii* infection in two bat cell lines, PaLu and PaKi, derived from lung and kidney tissues, respectively. Cells were infected with the highly virulent type I RH strain of *T. gondii* expressing mCherry, enabling direct visualization and quantitative assessment of infection at days one and three post-infection (Fig.1). Quantitative analysis revealed that RH parasites efficiently infected both PaLu and PaKi cells, with a substantial proportion of mCherry-positive cells detected as early as day 1 (Fig. 1).

**Figure 1.**
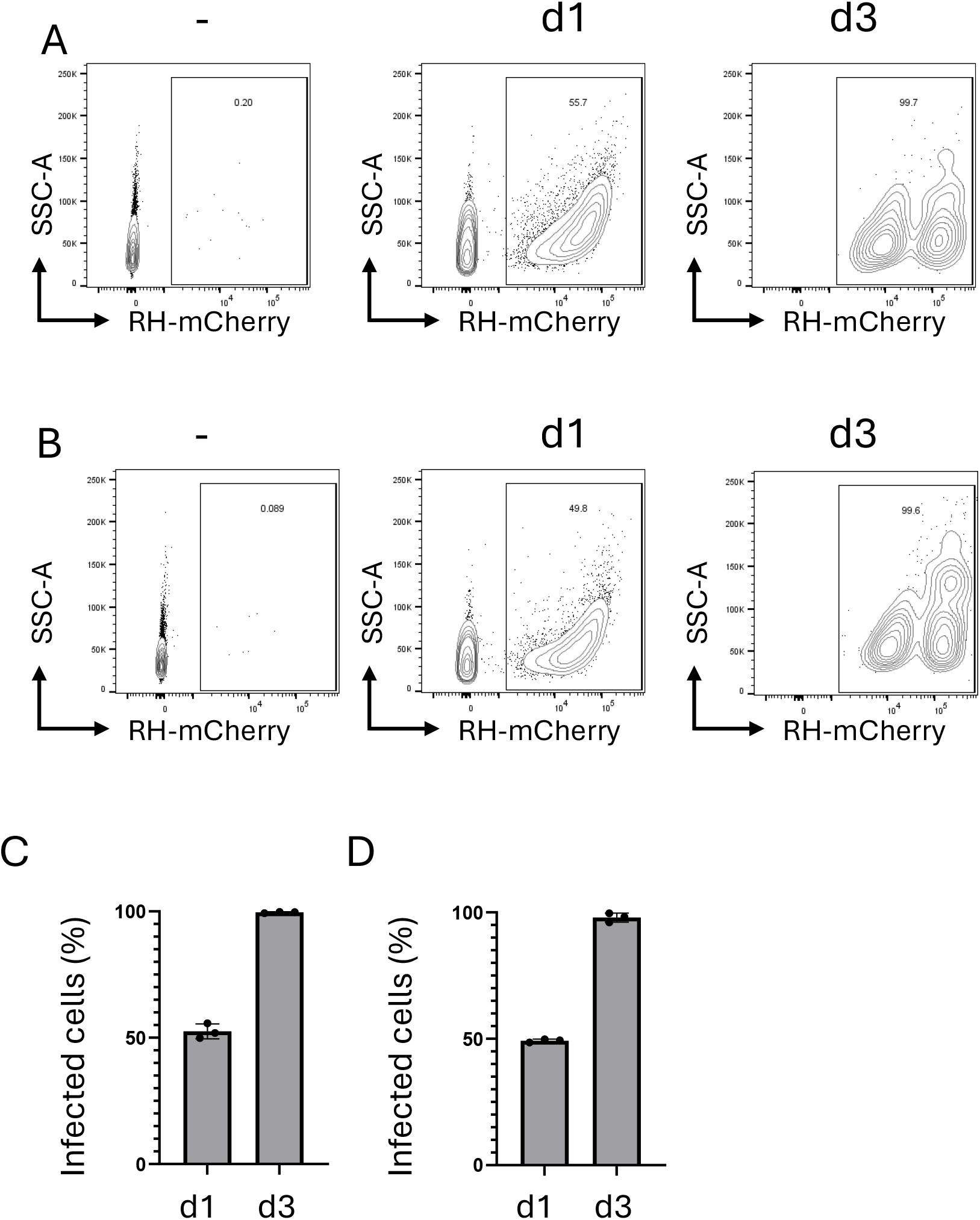
Bat epithelial cells are highly permissive to infection with the virulent type I RH strain of *T. gondii*. (A, C) PaLu and (B, D) PaKi cells were infected with the mCherry-expressing type I RH strain of *T. gondii* at a multiplicity of infection (MOI) of 3. The percentage of infected cells was determined by flow cytometry based on mCherry fluorescence at days 1 and 3 post-infection. Data are representative of four independent experiments, each performed with three independent cell cultures.

Infection progressed rapidly, and by day 3, nearly the entire cell population was infected in both cell lines. These findings demonstrate that bat epithelial cells are highly permissive to *T. gondii* infection and support efficient parasite propagation, establishing PaLu and PaKi cells as suitable experimental models for investigating bat cell-intrinsic responses to intracellular protozoan infection.

The RH strain represents a highly virulent type I lineage of *T. gondii*, in which infection with even a single parasite can be sufficient to cause lethality in susceptible mice [18-20]. To determine whether the high permissiveness of bat epithelial cells was limited to this highly virulent strain, we next examined infection with the type II strain Pru in the same PaLu and PaKi cell lines (Fig. 2). Flow cytometric analysis demonstrated that, similar to RH parasites (Fig. 1), Pru parasites readily invaded both bat cell lines and established productive infection (Fig. 2).

**Figure 2.**
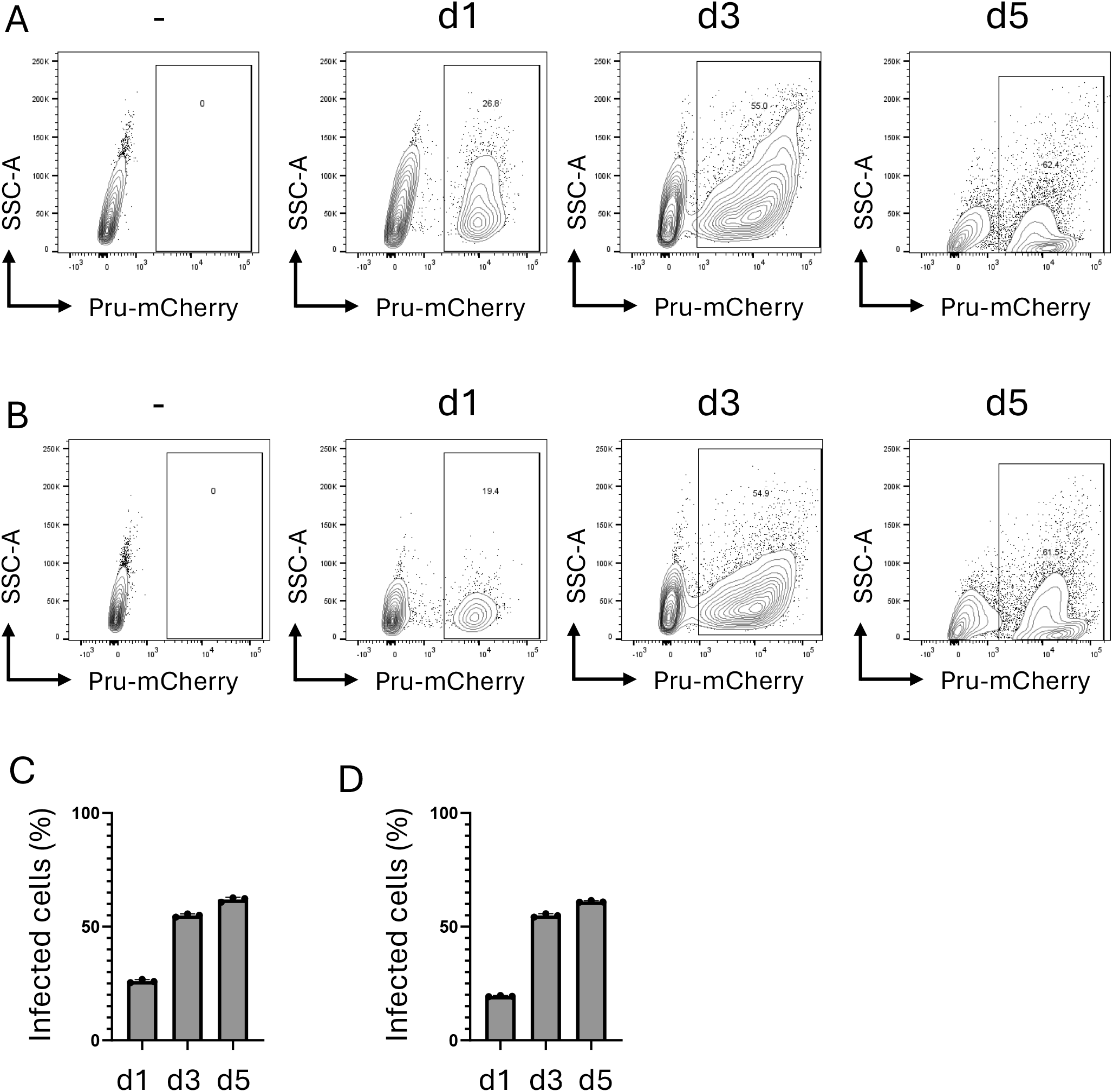
The type II Pru strain of *T. gondii* efficiently infects bat epithelial cells in vitro. (A, C) PaLu and (B, d) PaKi cells were infected with the mCherry-expressing type II Pru strain of *T. gondii* at an MOI of 3. The percentage of infected cells was determined by flow cytometry based on mCherry fluorescence at days 1, 3, and 5 post-infection. Data are representative of four independent experiments, each performed with three independent cell cultures.

However, the progression of Pru infection was slower than that observed with the RH strain (Fig. 1 and 2). Nevertheless, the proportion of infected cells increased over time, demonstrating that bat cell lines support infection by genetically and biologically distinct *T. gondii* strains. Together, these results establish that both type I RH and type II Pru parasites can efficiently infect bat cells and sustain successful infection *in vitro*.

### Bat epithelial cells display limited early responses to intracellular *T. gondii* infection

To characterize the transcriptional responses of bat cells to *T. gondii* infection, PaLu and PaKi cells were infected with the type II Pru strain and subjected to RNA sequencing at days 1, 3, and 5 post-infection. We selected the Pru strain rather than the highly virulent type I RH strain because RH infection resulted in rapid destruction of bat epithelial cell cultures, precluding a meaningful kinetic analysis of host transcriptional responses (Fig. 1).

Principal component analysis (PCA) revealed that both PaLu and PaKi cells undergo substantial transcriptional changes over time, even in the absence of infection (Fig. 3A and 3B). Indeed, uninfected epithelial cells collected on days 1, 3, and 5 exhibited distinct transcriptional profiles, indicating that bat cells actively respond to changes in their extracellular environment during culture (Fig. 3A and 3B). Remarkably, these culture-associated transcriptional changes remained a major source of variation even in *T. gondii*-infected cells, suggesting that environmental adaptation exerts a stronger influence on the global transcriptional landscape than intracellular parasite infection during the early stages of infection (Fig. 3 and SF1, SF2). Consistent with this interpretation, infected and uninfected samples showed limited separation at days 1 and 3, despite readily detectable intracellular parasite replication by day 3. Only at day 5 did infection-associated transcriptional changes become clearly apparent (Fig. 3A).

**Figure 3.**
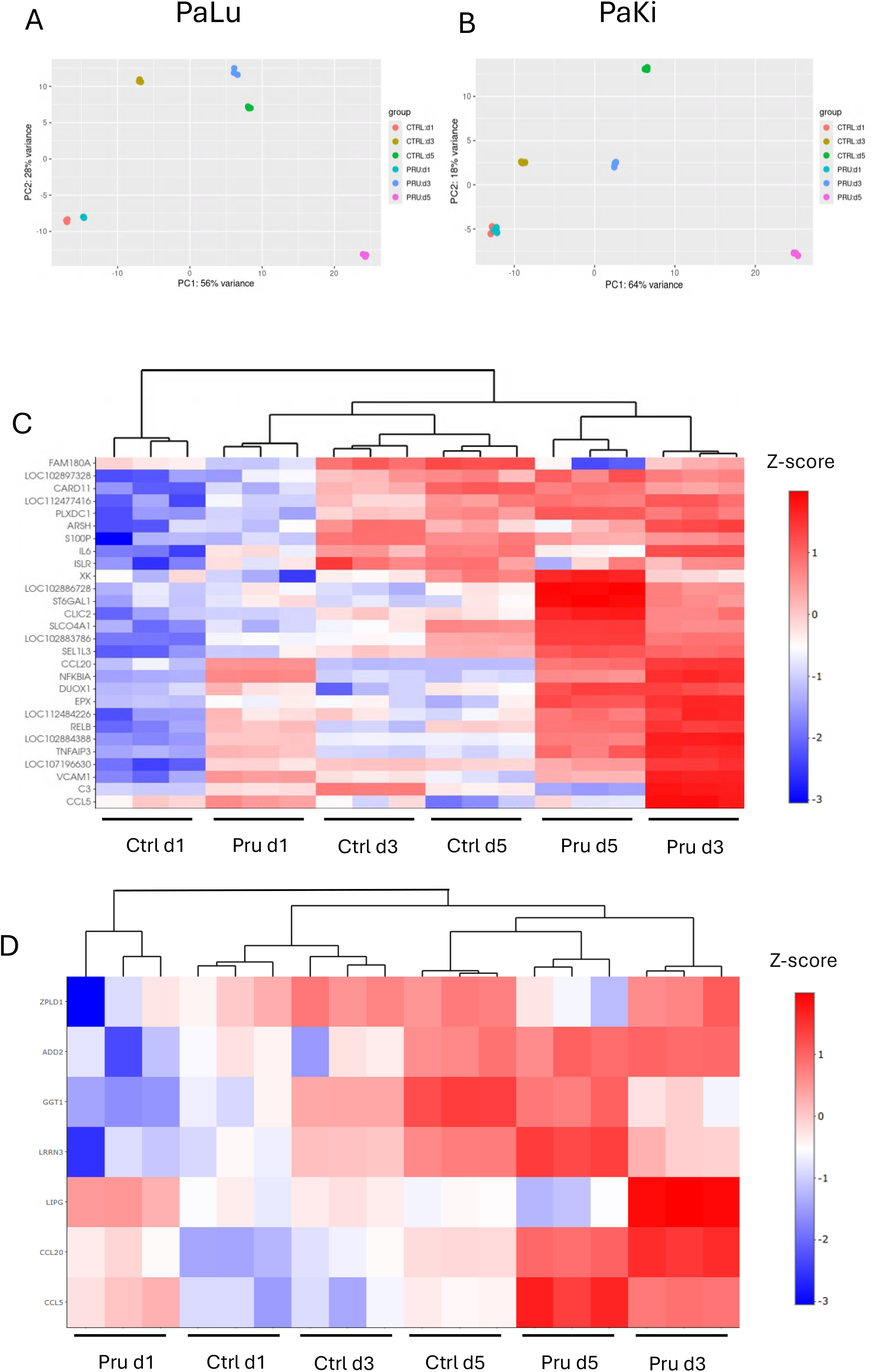
Transcriptional responses of bat epithelial cells to *T. gondii* infection. PaLu and PaKi cells were infected with the type II Pru strain of *T. gondii* at an MOI of 3, and total RNA was collected at days 1, 3, and 5 post-infection for RNA sequencing. Uninfected cells maintained under identical culture conditions were collected at the corresponding time points. (A, B) Principal component analysis (PCA) of gene expression profiles from infected and uninfected (A) PaLu and (B0 PaKi cells across the indicated time points. Each dot represents an individual sample, with colors indicating the six experimental groups: uninfected controls and Pru-infected cells collected on days 1, 3, and 5. PCA reveals substantial transcriptional variation associated with the duration of cell culture, whereas infection-associated differences become more apparent at day 5. (C) Heatmap illustrating gene expression patterns in PaLu cells across infection conditions and time points. (D) Heatmap illustrating gene expression patterns in PaKi cells across infection conditions and time points. The heatmaps illustrate temporal changes in gene expression and infection-associated transcriptional responses, highlighting the predominance of culture-associated transcriptional variation during the early stages of infection and the emergence of more pronounced infection-associated responses at day 5.

This unexpectedly delayed response suggests that bat epithelial cells can sustain substantial intracellular *T. gondii* replication (Fig. 1 and 2) without mounting a robust transcriptional response typically associated with pathogen recognition, inflammation, or cellular stress (Fig. 3). However, by day 5, when extensive parasite replication was accompanied by substantial cell damage and loss of cell viability, both PaLu and PaKi cells exhibited pronounced changes in gene expression (Fig. 3, SF1, SF2).

Analysis of the most highly upregulated genes revealed both shared and cell line-specific features of this late response. In PaLu cells, infection was associated with increased expression of genes involved in inflammatory signaling and tissue remodeling, including CCL20, SPP1, and LIPG, together with genes implicated in cellular metabolism and adaptation to stress, such as PDK4, GPR68, and HMGA1. Increased expression of ST6GAL1, MAL2, and SPINK1 further suggested alterations in glycosylation, membrane-associated processes, and cellular protective responses [21-27]. Collectively, these changes are consistent with a broad cellular adaptation program involving metabolic reprogramming, inflammatory mediator production, and tissue remodeling rather than a narrowly defined pathogen-specific immune response (Fig. 3, SF1).

PaKi exhibited a partially distinct transcriptional response, characterized by increased expression of CCL5, SEMA7A, and TNFRSF11B, genes associated with inflammatory signaling and cell– cell communication. Upregulation of RGS1 and RGS2 suggested changes in G protein-coupled receptor signaling, whereas increased expression of MMP16 was consistent with extracellular matrix remodeling (Fig. 3, SF2). Interestingly, PDK4 was upregulated in both PaLu and PaKi cells, suggesting that metabolic adaptation may represent a shared component of the late response to infection. [28, 29]

Together, these findings demonstrate that the transcriptional response of *P. alecto* epithelial cells to intracellular *T. gondii* infection is remarkably delayed. During the first three days of infection, transcriptional changes associated with the culture environment predominate, despite ongoing intracellular parasite replication. A pronounced infection-associated transcriptional program emerges only at day 5, coinciding with extensive parasite replication and overt cellular damage. Notably, this late response includes genes involved in metabolism, inflammatory signaling, and tissue remodeling, suggesting that bat cells, at least those of *P. alecto*, respond to advanced infection through broad cellular adaptation rather than an immediate, robust inflammatory transcriptional program.

## Discussion

Our study provides evidence that the distinctive relationship between bats and intracellular pathogens extends beyond viral infections to the protozoan parasite *T. gondii*. Using two independently derived *P. alecto* cell lines, we found that bat cells are highly permissive to infection with genetically and biologically distinct *T. gondii* strains. Both type I RH and type II Pru parasites readily invaded PaLu and PaKi cells and underwent progressive replication, ultimately resulting in widespread infection of the cultures. Despite this substantial intracellular parasite burden, bat cells mounted remarkably limited transcriptional responses during the early stages of infection.

The ability of *T. gondii* to efficiently infect bat cells is consistent with epidemiological evidence demonstrating natural exposure of diverse bat species to this parasite. Serological surveys have detected *T. gondii*-specific antibodies in multiple bat species and geographic regions, while parasite DNA and, in some studies, viable parasites have been recovered from naturally infected bats [16, 17]. These observations establish that bats can acquire *T. gondii* infection in nature but provide little information about the cellular interactions between the parasite and its bat host. Our results directly demonstrate that *P. alecto* cells provide a permissive intracellular environment for parasite invasion and replication. Importantly, this phenotype was not restricted to the highly virulent type I RH strain. Although Pru infection progressed more slowly, type II parasites also efficiently established infection and progressively spread throughout both bat cell cultures. Thus, the ability of *T. gondii* to replicate in bat cells appears to reflect a fundamental compatibility between the parasite and the bat intracellular environment rather than an unusual property of the RH strain.

One of the most striking findings of our study was the remarkably limited transcriptional response of bat epithelial cells during the initial stages of infection. Even at three days after Pru infection, when intracellular parasites were readily detectable and infection was progressing through the cultures, relatively little alteration in the host transcriptional program was observed. Interestingly, this limited response contrasted with substantial transcriptional changes observed in uninfected cells over the same culture period, suggesting that bat cells actively respond to changes in their environment while remaining relatively unresponsive to intracellular parasite infection. Moreover, these culture-associated transcriptional changes remained a dominant feature of the transcriptional landscape even in infected cells. A pronounced infection-associated transcriptional response emerged only at day 5, coinciding with a high parasite burden, extensive infection of the cultures, and substantial parasite-associated cell damage. This late response included genes associated with inflammatory signaling, metabolic adaptation, and tissue remodeling, suggesting that it may reflect both responses to infection and secondary consequences of cellular injury. Thus, rather than responding proportionally to the presence or replication of an intracellular pathogen, PaLu and PaKi cells appeared capable of accommodating considerable parasite replication before mounting a strong transcriptional response.

This phenotype is particularly intriguing in the context of the emerging concept of disease tolerance in bats. Much of the current understanding of bat immunity has developed from studies of viral infections, which suggest that bats combine effective antiviral mechanisms with mechanisms that limit potentially damaging inflammatory responses [1, 2]. Multiple features of bat immunity including alterations in DNA sensing, inflammasome pathways, interferon regulation, and inflammatory signaling have been proposed to contribute to this phenotype [4-6]. For example, the PYHIN family of cytosolic DNA sensors, which includes AIM2 and IFI16 in other mammals, is absent from several characterized bat genomes, including *P. alecto* [30, 31], while other innate immune pathways display evidence of evolutionary adaptation. Our results raise the possibility that reduced responsiveness to intracellular perturbation is not exclusively an antiviral adaptation but may represent a more general property of bat cells that becomes evident during infection with evolutionarily unrelated intracellular pathogens.

Importantly, however, our findings also illustrate a potential cost associated with such a strategy. A muted early host response may limit inflammation and cellular damage when a pathogen can otherwise be controlled, but in the case of *T. gondii*, the absence of effective parasite restriction permitted continued replication. By day 5, the transcriptionally quiescent early phase was replaced by a strong host response coincident with extensive parasite burden and cell death.

Thus, in this *in vitro* system, limited early responsiveness did not equate to resistance. Instead, bat epithelial cells appeared to accommodate intracellular parasite replication until the parasite burden itself became incompatible with cellular survival. This distinction between resistance, which reduces pathogen burden, and tolerance, and in turn limits the consequences of infection without necessarily reducing pathogen burden, may be particularly important for understanding bat–pathogen interactions [2].

Several limitations of the present study should be considered. First, our experiments were performed using immortalized cell lines from a single bat species. Bat immunity is highly diverse across Chiroptera, and responses observed in *P. alecto* cannot necessarily be generalized to all bat species. Second, epithelial cells provide a useful model for investigating cell-intrinsic immunity but do not reproduce interactions among immune and nonimmune cells that occur during infection *in vivo*. Third, the late transcriptional response observed in heavily infected cultures is likely to contain both infection-specific responses and secondary responses to cellular injury. Future studies using single-cell approaches, additional early sampling, and separation of infected from bystander cells should help distinguish these components.

Despite these limitations, our findings establish a framework for investigating antiparasitic immunity in bats and reveal potentially important differences from established mouse and human models of toxoplasmosis. *P. alecto* cells are highly permissive to *T. gondii* and exhibit surprisingly limited transcriptional responses during the early stages of intracellular infection, despite their capacity to undergo substantial transcriptional changes in response to culture-associated environmental conditions. These observations extend the concept of distinctive bat immune regulation beyond viral infection and suggest that cellular tolerance to intracellular pathogens may represent a broader feature of bat biology. At the same time, the extensive parasite replication ultimately observed *in vitro* emphasizes that tolerance and resistance are separable components of host defense. Defining the molecular pathways that allow bat cells to accommodate intracellular infection while limiting early inflammatory activation and determining how these pathways operate *in vivo* may provide important insights into the evolution of mammalian immunity and mechanisms that balance antimicrobial defense with protection from immunopathology.

## Methods

### Cell culture and T. gondii infection

The Pteropus alecto lung (PaLu) and kidney (PaKi) epithelial cell lines were maintained in RPMI 1640 medium supplemented with 10% fetal bovine serum (FBS) at 37°C in a humidified atmosphere containing 5% CO_2_. The mCherry-expressing type I RH and type II Pru strains of *T. gondii* were maintained in vitro by serial passage in human foreskin fibroblasts (HFFs). For infection experiments, PaLu and PaKi cells were inoculated with freshly harvested tachyzoites of either the RH or Pru strain at a parasite-to-host cell ratio of 3:1 (multiplicity of infection of 3).

Infected cells were maintained under the same culture conditions and analyzed at the indicated time points post-infection. The percentage of infected cells was determined by flow cytometry using a BD LSRFortessa flow cytometer (BD Biosciences) based on the detection of parasite-expressed mCherry fluorescence.

RNA Sequencing and Bioinformatics Analysis

Total RNA was isolated from uninfected and *T. gondii*-infected PaLu and PaKi cells at the indicated time points using a Qiagen RNA extraction kit according to the manufacturer’s instructions. RNA library preparation and high-throughput sequencing were performed by the University of Rochester Medical Center Genomics Research Center (Rochester, NY), using an Illumina NovaSeq X Plus sequencing platform with 150-bp paired-end reads.

Raw sequencing reads were demultiplexed from Illumina base call files using bcl-convert (v4.1.7). RNA-seq data were processed using the nf-core/rnaseq pipeline (v3.22.2), with reproducible software environments provided by the Bioconda and BioContainers projects [32-34]. Pipeline execution was managed using Nextflow (v25.10.0) [35]. Read quality control, adapter trimming, and filtering were performed using fastp (v0.24.0) [36]. Processed reads were aligned to the Pteropus alecto reference genome (ASM32557v1) using STAR (v2.7.11b) [37, 38]. Transcript abundance was quantified using Salmon (v1.10.3), and transcript-level estimates were aggregated to obtain gene-level expression estimates [39]. BedGraph files were generated using BEDTools (v2.31.1) and converted to BigWig format using the UCSC bedGraphToBigWig utility (v469).

Gene expression normalization and differential expression analysis were performed using DESeq2 (v1.52.0) in R (v4.6.1). Genes with a Benjamini–Hochberg-adjusted P value < 0.05 were considered significantly differentially expressed. Principal component analysis (PCA) was performed using DESeq2 to assess variation in gene expression among experimental groups [40]. Heatmaps were generated using the gplots (v3.3.0) and heatmaply (v1.6.0) packages with regularized log-transformed gene counts as input. Differentially expressed genes were visualized using volcano plots generated with ggplot2 (v4.0.3), with significantly upregulated and downregulated genes (adjusted P < 0.05) indicated in red and blue, respectively. Gene Ontology enrichment analysis was performed separately for upregulated and downregulated genes using Enrichr (v3.4). The most significantly enriched Gene Ontology terms were visualized using dot plots generated with ggplot2 (v3.5.2) .

## Figure legends

Supplemental Figure 1. Transcriptional responses of bat Palu cells to *T. gondii* infection.

Volcano plots showing differential gene expression in uninfected PaLu cells across time points from day 1 to day 5. Red dots indicate genes with an adjusted P value ≤ 0.05.

Supplemental Figure 2. Transcriptional responses of bat PaKi cells to *T. gondii* infection.

Volcano plots showing differential gene expression in uninfected PaKi cells across time points from day 1 to day 5. Red dots indicate genes with an adjusted P value ≤ 0.05.

## Acknowledgments

We are grateful to Ruth Serra-Moreno and Sydney Kei Brown for their assistance with bat epithelial cell culture, insightful discussions, and careful review of our manuscript.

## Contributions

P.Y. designed and conceived the research. P.Y., S.F.K., performed experiments. S.F.K. supervised the research. P.Y. analyzed data. P.Y. wrote the manuscript.

## Notes

### Competing Interest Statement

The authors have declared no competing interest.

