## Supplemental Figures for "Transcriptomic Analysis of Innate Immune Responses to *Toxoplasma gondii* in Bat Cells"

### Supplemental Figure 1

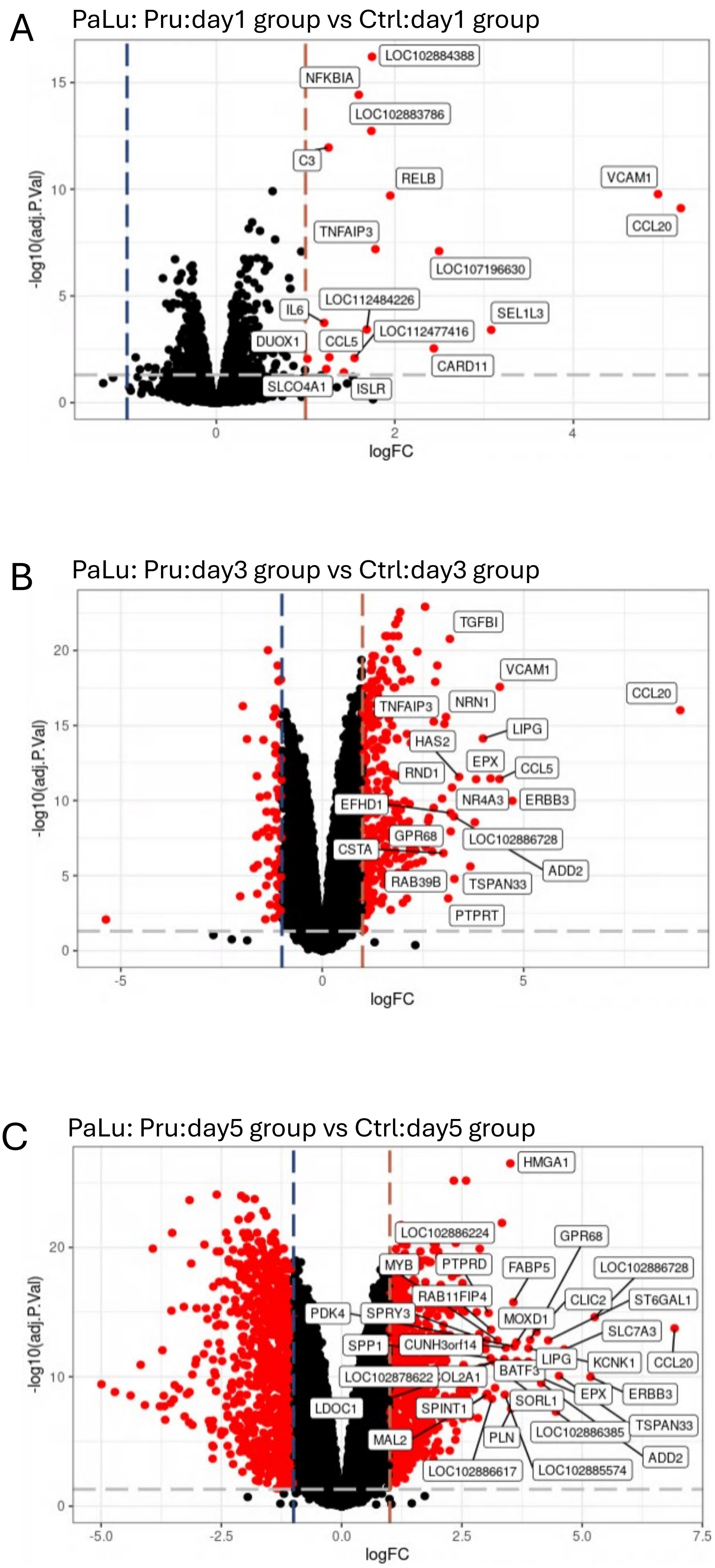

### Supplemental Figure 2

A

PaKi: Pru:day1 group vs Ctrl:day1 group

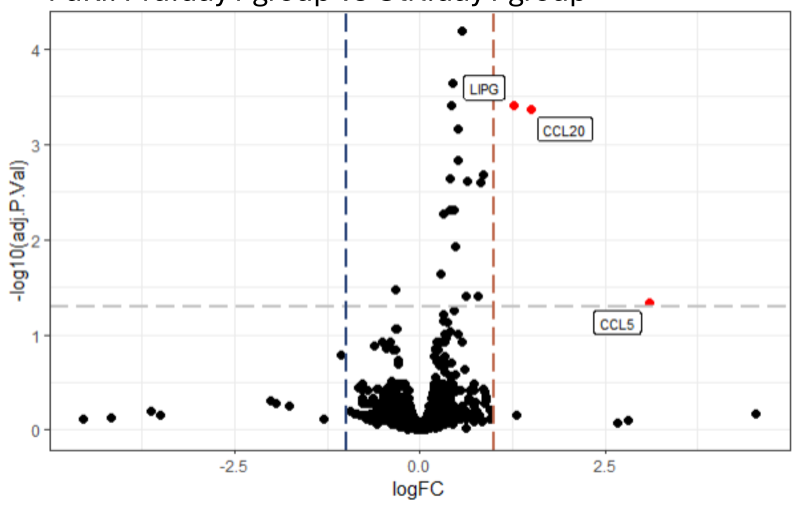


B

PaKi: Pru:day3 group vs Ctrl:day3 group

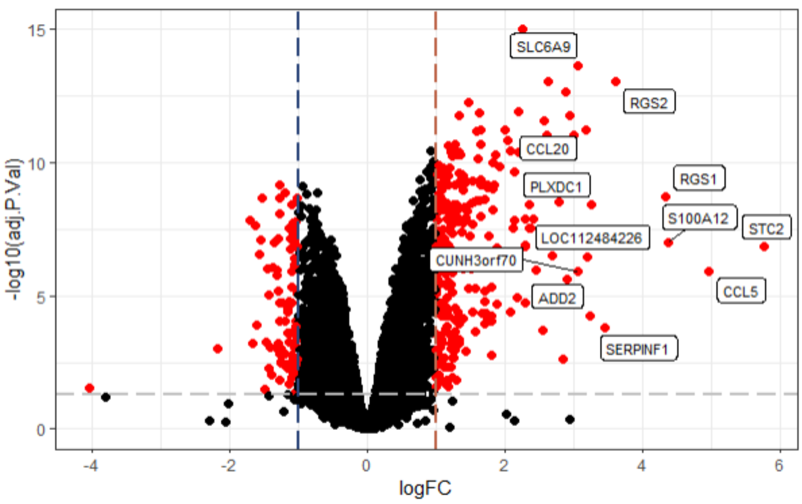


C

PaKi: Pru:day5 group vs Ctrl:day5 group

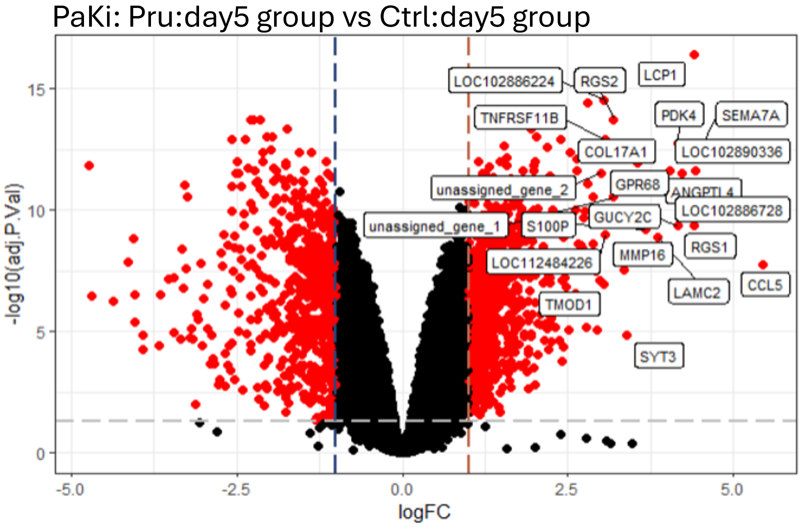
